# Population model of Temnothorax albipennis as a distributed dynamical system II: secret of “chemical reaction” in collective house-hunting in ant colonies is unveiled by operator methods

**DOI:** 10.1101/2021.07.14.452425

**Authors:** Siwei Qiu

## Abstract

The collective intelligence of animal groups is a complex algorithm for computer scientist and a many-body problem for physics of living system. We show how the time evolution of features in such a system, like number of ants in particular state for colonies, can be mapped to many-body problems in non-equilibrium statistical mechanics. There exist role transitions of active and passive ant between distributed functions, including exploration, assessing, recruiting and transportation in the house-hunting process. Theoretically, such a process can be approximately described as birth-death process where large number of particles living in the Fock space and particles of one sub-type transfer to a different sub-type with some probability. Started from the master equation with constrain of the quorum criterion, we express the evolution operator as a functional integral mapping from operators acting on Fock space in number representation to functional space in coherent state representation. We then read out the action from the evolution operator, and we use least action principal equations of motion, which are the number field equations. The equations we get are couple ordinary differential equations, which can faithfully describe the original master equation, and hence fully describe the system. This method provides us differential equation-based algorithm, which allow us explore parameter space with respect to more complicated agent-based algorithm. The algorithm also allows exploring stochastic process with memory in a Markovian way, which provide testable prediction on collective decision making.

## I. Introduction

The secret of “chemical reaction” within ant colony when they are collectively looking for new house since their old nest was destroyed can be very interesting problem that gathering interest from both multi-disciplines, including biology [1], physics [2], and computer science [3]. In the previous work of this series of papers, we discussed the role of quorum sensing and the reduced theory related to the quorum sensing. However, the theory we discussed are simply deterministic ordinary differential equations, and we did not discuss the probability theory basis. In this paper, we build the link between the deterministic ODE and the reaction equations, making use of concept of Fock space in theoretical physics. Without making use of probability theory, we simply use operator method [4] to construct the path integral of the system, from which we derive equation of motion. In previous work, we made use of the complexity of the information in environment and taking both ant and their perceptive cues in environment. In this work, we step back and use simple constrain of quorum rule and focus on understanding the way we derive equation of motion. Notice that this method allows derivation of theory beyond mean field. To keep this work simple, we simply focus on derivation of mean field equation. In principle, same method can be used in network including spatial dependence, but we leave the extension to future work. In section II, we will discuss the mathematical derivation of equation of motion. In section III, we briefly show the numerical simulation result of the system using derived equations.

## II. Gap between mean field theory and original system

Let us first refresh some definitions related to our theory.

### Definition 1

Let *H* be a complex Banach space. Then *H* is called Hilbert space if a complex number 〈*x, y*〉 is the inner product of complex vector *x* and *y*, and such an inner product meet the following conditions:

1. 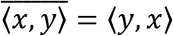
2. 〈*αx* + *βy, z*〉 = *α*〈*x, z*〉 + *β*〈*y, z*〉 for any scalar *α* and *β*
3. 〈*x, x*〉 = ‖*x*‖_2_

The definition above allows us to define the state of ant hunting model using a vector space, we then introduce “quantum” notation of vector using bra 〈*ϕ*| and ket |*ϕ*〉. So |*ϕ*〉 is the state vector, while 〈*ϕ*| is the dual vector, usually defined as complex conjugate transpose of the vector |*ϕ*〉.

In this article, we try to extend the definition of Hilbert space to the integrable function space. Due to the fact that we are interested in the change of ant number, our state function has discrete property, which extend the domain.

### Definition 2

Let *F* be a Fock space. The |*f*〉 denote a vector on Fock space, and 〈*f*| is in the dual vector. |*f*〉 is expended by basis |*n*〉,, while 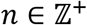. For each *n*, it is a tensor product of n Hilbert space. And Fock space is direct sum of all the |*n*〉. The inner product is defined as:

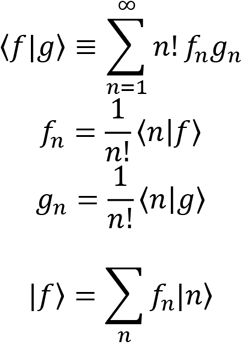

And this makes sense, with a complete base relation ∑*_n_* |*n*〉〈*n*| = 1.

If there are multiple sub-type of particle, like 3 of them, one can denote as |*n*_1_, *n*_2_, *n*_3_〉, representing the state of the mixture of 3 sub-type of particles have configuration (*n*_1_, *n*_2_, *n*_3_). For convenient, we will write such a composite state as 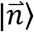, where 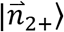 mean |*n*_1_, *n*_2_ + 1, *n*_3_〉 and 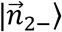 mean |*n*_1_, *n*_2_ – 1, *n*_3_〉.

### Theorem 1

With the following definition,

i. |*ϕ*〉 = ∑*_n_ϕ_n_*|*n*〉, |*ψ*〉 = ∑*_n_ψ_n_*|*n*〉 are two state vectors in generalized Hilbert space.
ii. *ϕ*(*z*) = *ϕ_n_z^n^* and *ψ*(*z*) = *ψ_n_z^n^* are the generating function

We then can have the following integral form of inner product:

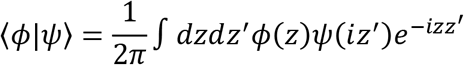

The integral for *z* and *z*′ both run through real line.

Proof: The inner product can be written as follows:

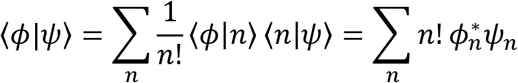

We then make use of the following identity to simplify the expression:

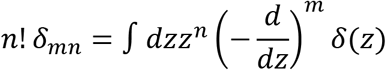

We then get:

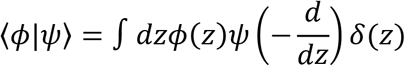

We also know:

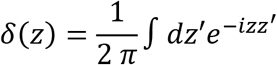

We see that:

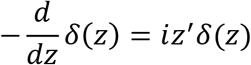

This means we can replace 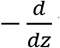 with *iz*′. Actually, later we will see that this is the property of annihilation operator, where *iz*′ is the eigenvalue of the operator, and *δ*(*z*) here is the eigenfunction, which will have a vector counterpart in vector space.

Based on this, we can write down our desired relation:

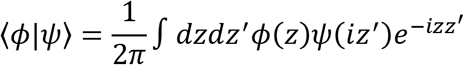

Up to here, we already get some flavor of the theory, we see that the role of operator 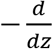 is making transition from order *n* to order *n* – 1, which can be easily seen by using the generating function *ϕ*(*z*) = *ϕ_n_z^n^*, where 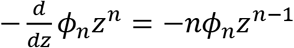. However, when look at operator z, it does opposite, since *zϕ_n_z^n^* = *ϕ_n_z*^*n*+1^. For physicist, we know that this is same as the creation and annihilation operator *a*^†^ and *a*. We can easily also check that there is commutation relation [*a*, *a*^†^] ≡ *aa*^†^ – *a*^†^*a* = 1, which fit to operator *z* and 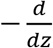. We thus can define algebra of operators based on the generating functions and create field theory using the eigenvalue of these operators. This explains why the classic problem can correspond to the bosonic quantum theory. We will show that this allows us to translate the master equation into differential equation, which simplify our biology problem.

### Definition 3

We are interested in dynamical system where the state is evolving as follows:

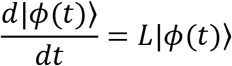

We then have the evolution operator:

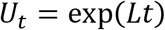

The ansatz of the state evolution is:

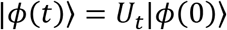

### Theorem 2

We have the path integral form of evolution operator *U_t_*:

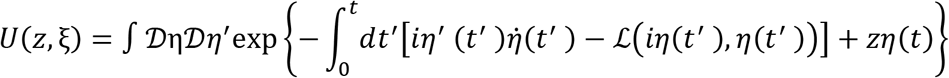

Where the boundary condition is:

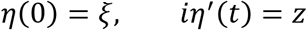

The evolution operator can be written in the following form:

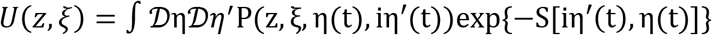

And we can get the mean field theory equation by imposing least action principal:

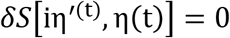

For the ant house hunting model, we consider the following 4 transitions, with *M* nests:

1. We denote 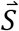 as the searching population. All searching ant has initial home nest as nest 0, which has quality 0.
2. We denote 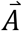 as the assessment population. Transition from searching state to assessment state can also cause by ant with home ID *i*. get recruited by the recruit ant doing tandem run for site *j*, and follow that ant to site *j*. This can happen in nest *k* if this lead ant for site *j* has home in nest *k*. Since search ant have equal possibility to be in any nest, we only need to take nest *i*. and nest *j* into account.
3. We denote 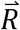 as the recruitment population. For transition from assessment ant for nest *i*. with old home nest index as *k* to the recruiter in tandem run for nest *i* has a transition rate *κ_i_*. At the same time, for simplicity, this rate is the average rate for old home nest index from 1 to *M*, where the value for each possible old home is *g*(*q_k_* – *q_i_*), where *g*(*x*) is a logistic function with form 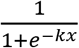. The transition rate should be 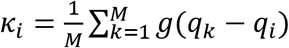.
4. We denote 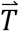 as the transport population, which include both passive ant and brood items. We assume the transport speed is proportional to Φ_i_, which is function of recruit population *R_i_*. Intuition is that if *R_i_* > *Th*, where *Th* is the threshold number of recruiters, then Φ_i_ become some positive value. However, before recruiter number reach the threshold, Φ_i_ remain zero.

In order to solve the ant problem, we write down the master equations as follows:

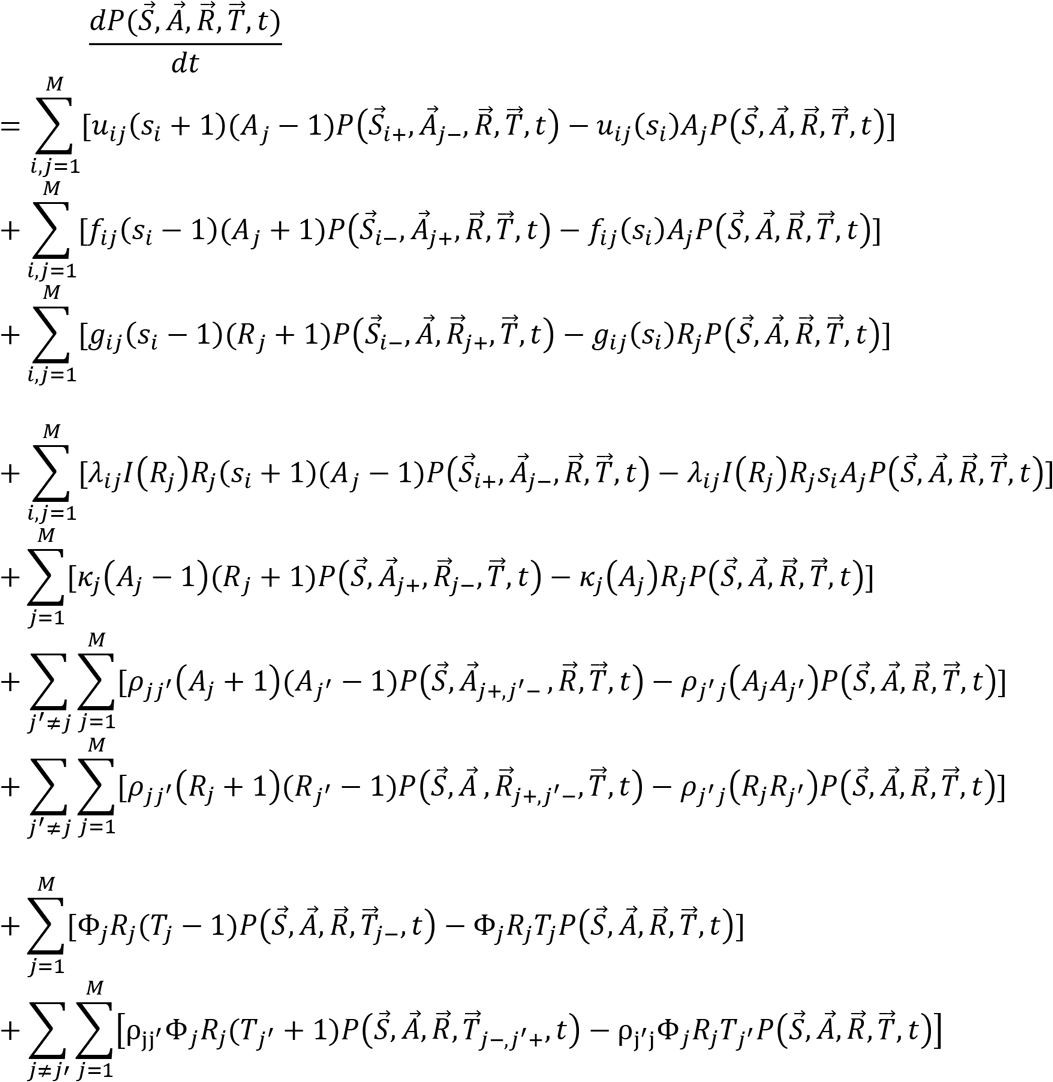

We assume there are *M* ant nests. Function *I*(*R, S*) is one if *R* is less than threshold, but zero if *R* is larger.

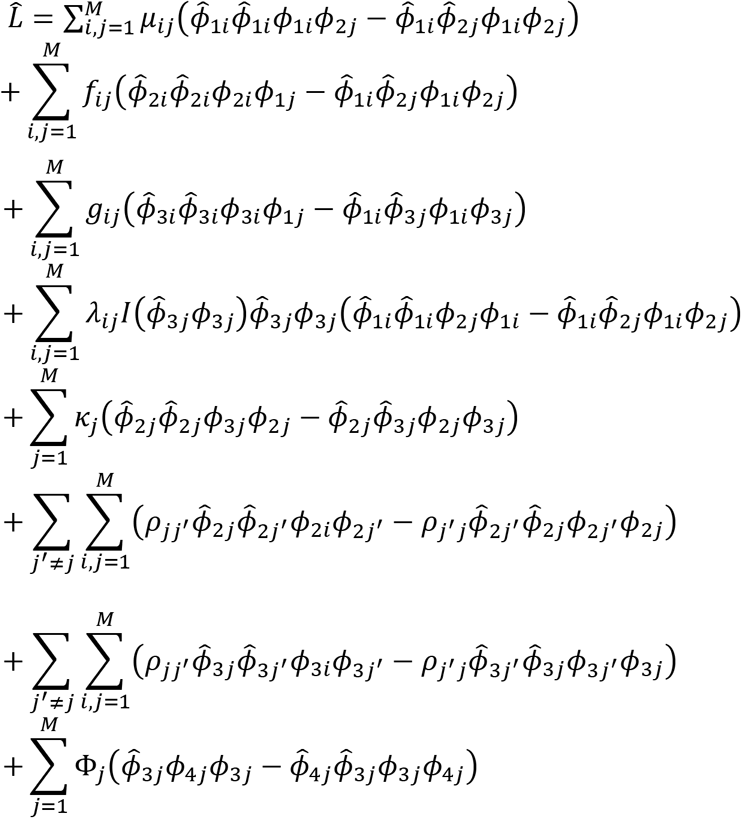

We denote the eigenvalue of operator *ϕ* to be *φ*, and the eigenvalue of dual operator 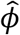 to be 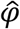, notice also we shift the field 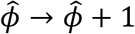 making use of the commutation relation of operator algebra.

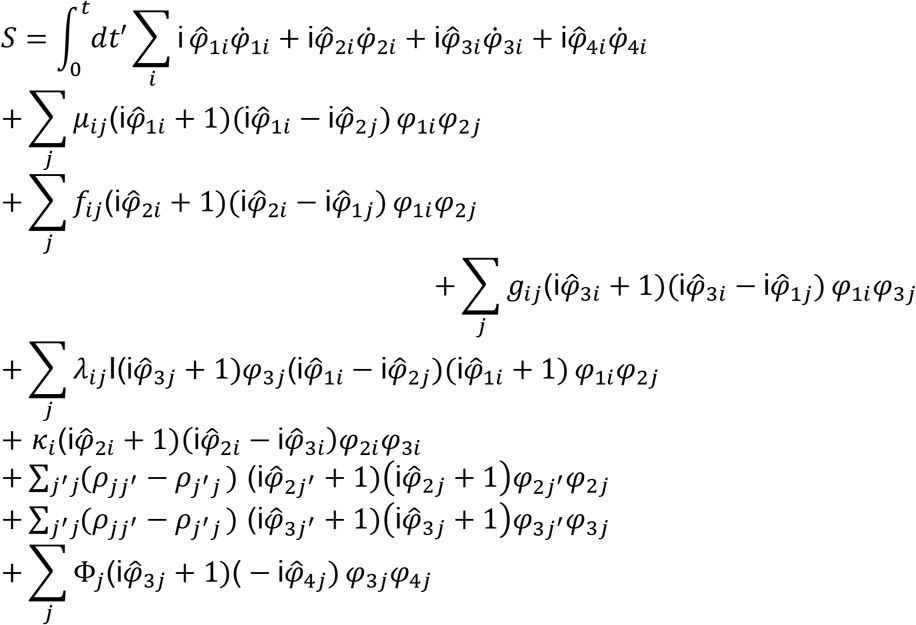

Up to first order, we have the following dynamical system:

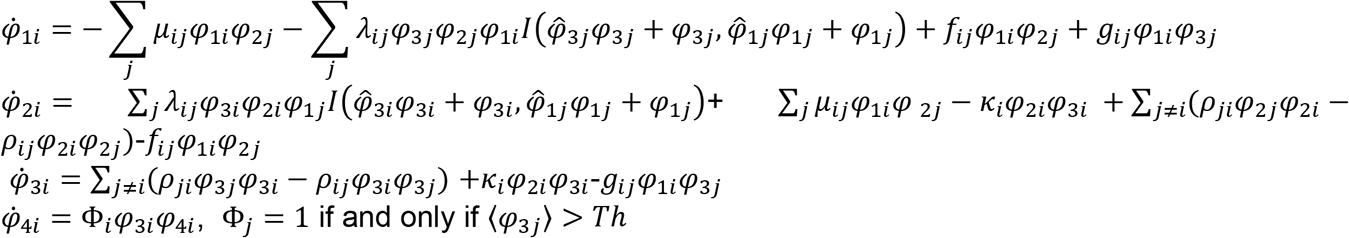

Remark: It is necessary to analyze the scale a bit. All the fields 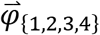 are representing the number of particles, so they are of order 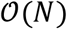. Here we assume total number of ants in whole system is *N*. One can imagine *μ_ij_*, *λ_ij_*, *κ_i_* and Φ*_i_* are of order 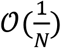, because quantity like *μ_ij_φ*_2*j*_ need to be a finite number. Similarly, we can assume the auxiliary fields 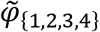 are all with order 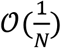. In this way, we can explain for eigenvalue of number operator, of form 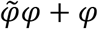, the leading term is *N* times larger than the second order term 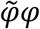, which is why we can write the equations of first order as follows:

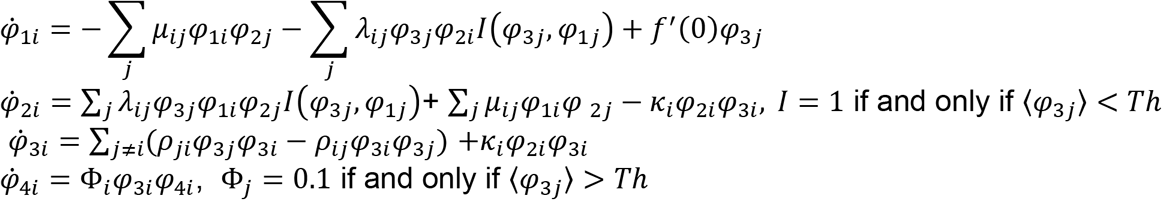

If we want to do higher order correction using the action, we need to keep the scale in mind, and we will know how big the correction should be based on the scale we know.

If we ignore the information transmission between recruiter, we have following equations:

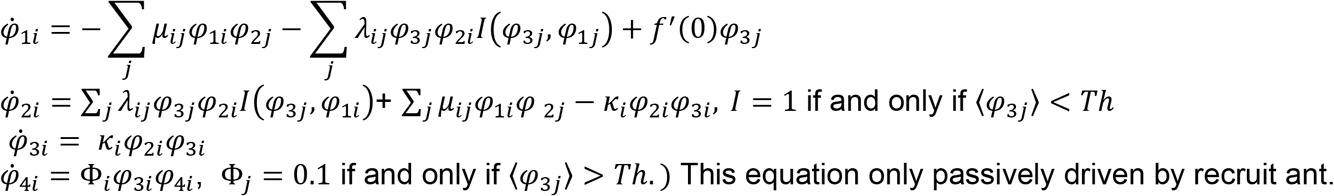

The term *f*′(0)_*φ*_3*j*__ is coming from the assumption based on Jiajia’s paper. If we do not consider this term, we have simple equations:

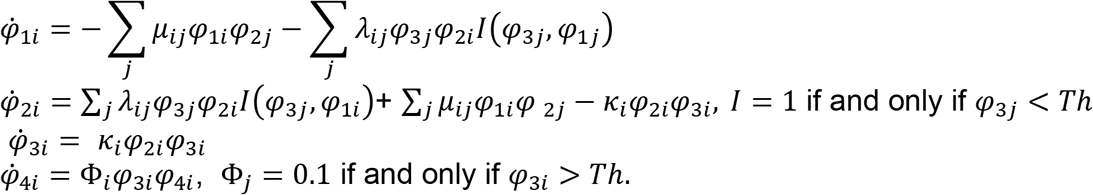

This is only passively driven by recruit ant.

These equations fully describe the distributed system of ant house-hunting, as long as the total ant number is large enough, the higher order correction only give small contribution. The solution of the 4^th^ equation is of this form:

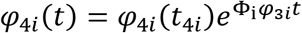

Where *t*_4*i*_ is the time when *φ*_3*i*_ is larger than the threshold *Th*. The smallest *t*_4*i*_ shows the winner. Converge rate should be roughly Φ_i_*Th* for the transportation, it is exponentially fast.

It is reasonable just simulate the first 3 equation sets. These are totally 9 equations for 3 nests.

## III Numerical simulation

Here we show some result of numerical simulation of the mean field equation just to show that the resulting equations describe the system’s performance well.

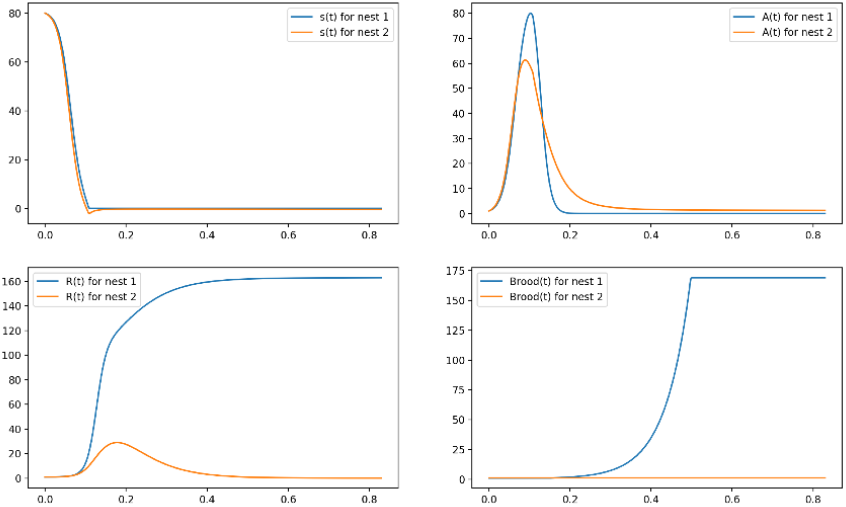
Fig Simulation result using derived mean field equation.

## IV Discussion

We revisit the method from Feyman’s path integral method [5–7] to derive a mean field equation that can reproduce the behavior of agent-based computational model [3]. This method also gives us the tool to explore the dynamics beyond mean field, one simply just need to linearize the system around stationary solution and then compute the propagator equations. The higher moment can be constructed through Feynman rule. These methods have been used in both physics and theoretical neuroscience literature. In the future, we will extend the system to include spatial dependency, meaning that the distance between nests will matter. This will introduce the competition between the quality of the nests and the effort to reach the nest, making the decision making more challenging.

## Notes

### Competing Interest Statement

The authors have declared no competing interest.

